# Differential auditory and visual phase-locking are observed during audio-visual benefit and silent lip-reading for speech perception

**DOI:** 10.1101/2021.12.18.472955

**Authors:** Máté Aller, Heidi Solberg Økland, Lucy J. MacGregor, Helen Blank, Matthew H. Davis

## Abstract

Speech perception in noisy environments is enhanced by seeing facial movements of communication partners. However, the neural mechanisms by which audio and visual speech are combined are not fully understood. We explore MEG phase locking to auditory and visual signals in MEG recordings from 14 human participants (6 females, 8 males) that reported words from single spoken sentences. We manipulated the acoustic clarity and visual speech signals such that critical speech information is present in auditory, visual or both modalities. MEG coherence analysis revealed that both auditory and visual speech envelopes (auditory amplitude modulations and lip aperture changes) were phase-locked to 2-6Hz brain responses in auditory and visual cortex, consistent with entrainment to syllable-rate components. Partial coherence analysis was used to separate neural responses to correlated audio-visual signals and showed non-zero phase locking to auditory envelope in occipital cortex during audio-visual (AV) speech. Furthermore, phase-locking to auditory signals in visual cortex was enhanced for AV speech compared to audio-only (AO) speech that was matched for intelligibility. Conversely, auditory regions of the superior temporal gyrus (STG) did not show above-chance partial coherence with visual speech signals during AV conditions, but did show partial coherence in VO conditions. Hence, visual speech enabled stronger phase locking to auditory signals in visual areas, whereas phase-locking of visual speech in auditory regions only occurred during silent lip-reading. Differences in these cross-modal interactions between auditory and visual speech signals are interpreted in line with cross-modal predictive mechanisms during speech perception.

**Significance Statement:** Verbal communication in noisy environments is challenging, especially for hearing-impaired individuals. Seeing facial movements of communication partners improves speech perception when auditory signals are degraded or absent. The neural mechanisms supporting lip-reading or audio-visual benefit are not fully understood. Using MEG recordings and partial coherence analysis we show that speech information is used differently in brain regions that respond to auditory and visual speech. While visual areas use visual speech to improve phase-locking to auditory speech signals, auditory areas do not show phase-locking to visual speech unless auditory speech is absent and visual speech is used to substitute for missing auditory signals. These findings highlight brain processes that combine visual and auditory signals to support speech understanding.

## Introduction

Speech is the most important form of human communication and conventionally used in face-to-face conversation. Many types of adverse listening conditions decrease speech intelligibility, such as listening to a speaker with a foreign accent, in the presence of background noise or with a hearing impairment (see Mattys et al., 2012 for a review). Seeing the face of the conversation partner benefits speech comprehension in all such adverse conditions, both in healthy people and those with hearing impairment (Sumby and Pollack, 1954; Erber, 1975; Summerfield et al., 1992). Yet, the neural mechanisms of this benefit are still not fully understood. Behavioural evidence suggests that mouth movements are the main carrier of visual speech information. For example, eye-tracking demonstrates that listeners fixate the mouth more often when speech is harder to understand (Yi et al., 2013) and selective masking of oral movements is detrimental to comprehension (Thomas and Jordan, 2004). Furthermore, people with better silent lip reading ability benefit more from visual speech when listening to audio-visual speech (MacLeod and Summerfield, 1987).

Extensive research indicates that neural activity in auditory cortex synchronizes, or entrains (in a broad sense, see Obleser and Kayser, 2019), to temporally regular stimuli (Lakatos et al., 2005). Importantly, several studies demonstrated neural entrainment to auditory speech signals including their temporal envelope (Giraud and Poeppel, 2012; Peelle and Davis, 2012; Gross et al., 2013; Peelle et al., 2013; Ding and Simon, 2014). Additionally, there is close temporal correspondence between the auditory (acoustic envelope) and visual (lip aperture area) components of speech (Chandrasekaran et al., 2009). The prediction follows, that neural responses also track visual speech signals. Indeed, neural responses entrain to speakers’ lip movements in various listening situations: clear audio-visual speech from a single speaker (Luo et al., 2010; Micheli et al., 2020; Mégevand et al., 2020), clear audio-visual speech from multiple speakers (Zion Golumbic et al., 2013; Park et al., 2016), silent visual-only speech (O’Sullivan et al., 2017; Hauswald et al., 2018; Bourguignon et al., 2020; Nidiffer et al., 2021), and audio-visual speech-in-noise (Keitel et al., 2018). Here, we investigated how visual and auditory speech signals are combined to support comprehension without segregation of speech and background noise. We used noise-vocoded speech (Shannon et al., 1995) which is a form of intrinsic speech degradation with reduced spectral detail similar to that conveyed to hearing impaired individuals using a cochlear implant. Understanding such degraded speech is challenging even when a single sound source is present and is improved by visual speech. We hypothesized a mechanism by which such improvement can occur whereby neural entrainment to visual speech facilitates neural phase locking to degraded auditory speech (Peelle and Sommers, 2015).

Participants listened to audio-visually presented sentences and repeated as many words as they could in each sentence. We factorially manipulated the acoustic clarity of the sentences (high vs. low) using noise vocoding and the availability of visual speech (present vs. absent). This enabled us to assess how neural phase-locking to speech changes in response to increased sensory detail emanating from different sensory modalities (auditory or visual). To measure neural phase-locking to speech signals we collected magnetoencephalography (MEG) recordings and computed their phase coherence with i) the acoustic envelope of the auditory speech signal and ii) the time course of the instantaneous area of the speakers’ lip aperture (visual speech envelope) extracted from the sentence stimuli.

We replicated previous results showing that auditory and visual coherence effects emerge predominantly in temporal and occipital areas, respectively. We go beyond these by using partial coherence analysis to assess entrainment to auditory envelope signals in visual cortex and vice-versa for visual envelope signals in auditory regions. Given the previously shown coherence of auditory and visual speech signals, partial coherence analysis allows tests for true cross-modal influences in which additional neural variance is explained by auditory signals over and above entrainment to the visual input (and vice-versa). We further explore the relationship between neural phase-locking and behavioural measures of: (1) word report for visual only speech (i.e., lip-reading ability), and (2) the difference between word report for low-clarity audio-visual and high-clarity audio-only speech (a measure of audio-visual benefit).

## Materials and Methods

### Participants

Seventeen right-handed participants took part in the study after giving informed written consent. One participant did not finish the experiment and data from two participants were excluded because: (1) they were subsequently revealed to not be a native speaker of English or (2) data showed excessive MEG artefacts. The remaining 14 participants (6 females, 8 males, mean age ± SD = 28 ± 7 years) were native speakers of British English and had no history of hearing impairment or neurological diseases based on self-report. The experiment was approved by the Cambridge Psychology Research Ethics Committee and was conducted in accordance with the Declaration of Helsinki.

### Experiment design

Participants watched and listened to video clips of a speaker producing single sentences. We manipulated acoustic clarity (high vs. low) using noise vocoding (see Stimuli and procedure) and the availability of visual speech (present or absent) in a 2 x 2 factorial design resulting in 4 conditions: audio-only high clarity (AO_high_), audio-only low clarity (AO_low_), audio-visual high clarity (AV_high_) and audio-visual low clarity (AV_low_). A fifth condition with silent visual-only speech (VO) was also included (Fig. 1A).

**Figure 1.**
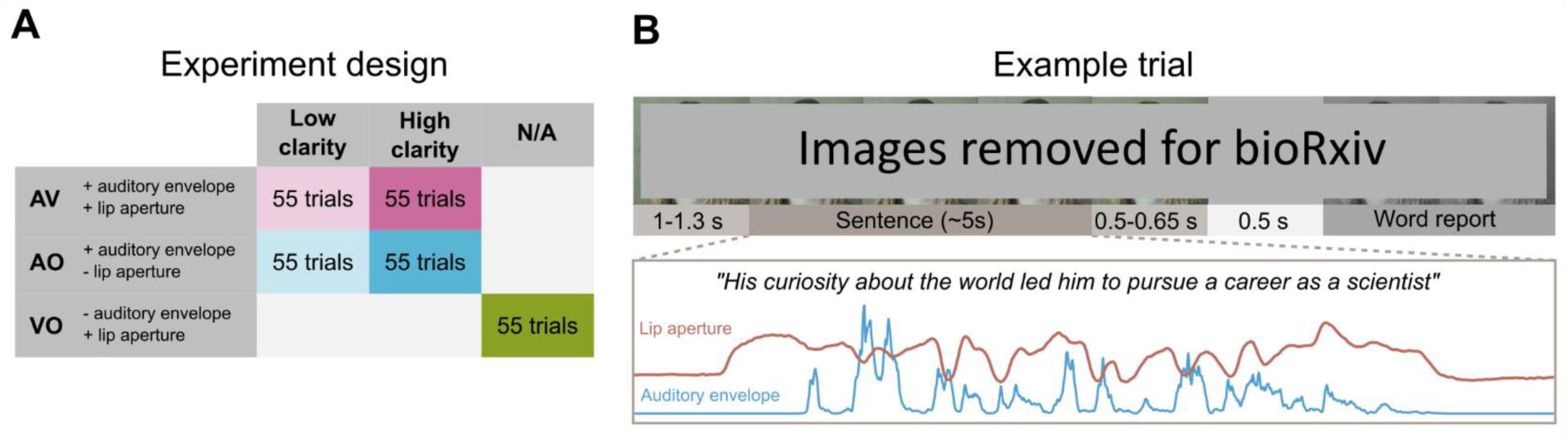
Experiment design and example trial. **A.** Experiment design. **B.** Trial diagram showing the order and timing of events in a representative trial (top). Time course of the extracted auditory (acoustic envelope) and visual speech signals (lip aperture area) for an example sentence (bottom).

### Stimuli and procedure

A total of 275 meaningful sentences were used ranging in length from 8 to 21 words (mean ± SD = 14.0 ± 1.9) and in duration from 3.72 to 7.02 s (mean ± SD = 5.13 ± 0.59). All were produced by a female native speaker of British English and recorded using a digital video camera (Panasonic AG-AF101 HD) and external microphone (RØDE NTG2 Shotgun). Video and audio were digitized at 48 kHz, 16 bit and edited using Adobe Premiere Pro CS6, Adobe Audition 3.0, Praat (https://www.fon.hum.uva.nl/praat/), and Matlab (MathWorks Inc.). The video clips depicted the speaker’s face in front of a neutral background (see Fig. 1B). Video clips of 55 different sentences were presented in each of the 5 conditions (AO_high_, AO_low_, AV_high_, AV_low_, and VO) in random order. Each of the sentences was presented once for each participant and the particular sentences assigned to each condition were randomized across participants.

The availability of visual speech was manipulated by either including the video of the speaker producing the actual sentences (visual speech present) or including a video of the face of the speaker while they were not speaking (visual speech absent). Acoustic clarity was manipulated using noise vocoding (Shannon et al., 1995), based on a protocol used in a previous experiment (Zoefel et al., 2020). Briefly, the speech signal was first filtered into 16 logarithmically spaced frequency bands between 70 and 5000 Hz, and the amplitude envelopes were extracted for each band (half-wave rectified, low-pass filtered below 30 Hz). The envelope for each of those frequency bands, 𝑒𝑛𝑣(𝑏), was then mixed with the broadband envelope, 𝑒𝑛𝑣(𝑏𝑟𝑜𝑎𝑑𝑏𝑎𝑛𝑑), of the same speech signal in proportion 𝑝, to yield an envelope for each frequency band 𝑒𝑛𝑣_𝑓𝑖𝑛𝑎𝑙_(𝑏).

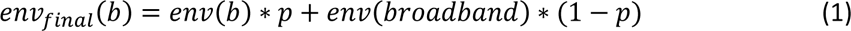

The envelopes were then used to modulate white noise in their respective frequency bands and the resulting signals were recombined. If 𝑝 = 0, then each of the narrowband envelopes become identical to the broadband envelope, hence the resulting signal is equivalent to 1-channel vocoded speech, which is unintelligible (Peelle et al., 2013). Conversely, if 𝑝 = 1, the resulting signal is equivalent to 16-channel vocoded speech which is fully intelligible (Peelle et al., 2013). This procedure enabled more precise control over acoustic clarity than achieved by changing the number of vocoder channels (see Zoefel et al., 2020 for another use of this method). In our experiment we used 𝑝 = 0.2 and 𝑝 = 0.7 for the low and high acoustic clarity conditions, respectively. The exact values for 𝑝 were determined in a separate pilot experiment (11 participants, 6 females, 5 males, mean age ± SD = 25 ± 2 years, recruited independently from the main experiment) such that: (1) listeners achieved approximately equal word report accuracy in AO_high_ and AV_low_ conditions and (2) the mean word report accuracy in these two conditions was at an intermediate value (close to 50% word report). In the visual-only condition the silent video of the speaker producing the sentence was presented.

Stimuli were delivered using Psychtoolbox version 3 (Brainard, 1997; Kleiner et al., 2007) running on Matlab 2014a. Each trial started with a fixation period where the non-speaking face of speaker was presented. After a delay of 1-1.3 s (uniformly sampled from 1, 1.1, 1.2 or 1.3 s) the sentence was presented, followed by a variable fixation period of 0.5-0.65 s (uniformly sampled from 0.5, 0.55, 0.6 and 0.65 s). Finally, participants were prompted by a brief (0.5 s) response cue to report verbally as many words as they could comprehend from the sentence or say “I don’t know” if they could not identify any words (Fig. 1B). The trial ended when participants pressed a button with their right hand indicating that they had finished speaking. Participants were instructed to fixate on the speaker’s face throughout the experiment. Participants received a short period of behavioural practice to familiarize themselves with vocoded speech at different levels of acoustic clarity. Sentence presentation was paired with a written transcription of each sentence to ensure efficient perceptual learning (Davis et al., 2005). They also practiced the word report task for vocoded speech before the main MEG experiment.

### Data acquisition and pre-processing

The verbal word report responses were audio recorded for off-line transcription. The transcription and word report accuracy scoring were done semi-automatically using custom python code. First, the verbal responses were processed with a speech recognition algorithm (python SpeechRecognition library; https://pypi.org/project/SpeechRecognition/) and the transcribed responses were manually checked. Then, for each sentence, the transcribed responses were compared to the corresponding original sentences using custom python code. The word report accuracy score was computed as the percentage of words correctly recognized irrespective of word order and averaged across sentences within each of the 5 conditions (AO_high_, AO_low_, AV_high_, AV_low_, and VO) separately for each participant.

Magnetic fields were recorded with a VectorView system (Elekta Neuromag) containing a magnetometer and two orthogonal planar gradiometers at each of 102 positions within a hemispheric array. Electric potentials were simultaneously recorded using 70 Ag/AgCl sensors according to the extended 10–10 system (EasyCap) and referenced to a sensor placed on the nose. All data were digitally sampled at 1 kHz and filtered between 0.03-330 Hz. Head position and EOG activity were continuously monitored using four head position indicator (HPI) coils and two bipolar electrodes, respectively. A 3D digitizer (Polhemus Fastrak) was used to record the positions of the EEG sensors, HPI coils, and ∼70 additional points evenly distributed over the scalp, relative to three anatomical fiducial points (the nasion and left and right pre-auricular points). Data from EEG sensors were not analysed further. Data from the MEG sensors (magnetometers and gradiometers) were processed using the temporal extension of Signal Source Separation (tSSS; Taulu and Simola, 2006) as implemented in Maxfilter 2.2 (Elekta Neuromag) to suppress noise sources, compensate for head movements, and interpolate any sensors that generated poor quality data. Finally, a band stop filter at 50 Hz was applied and the data were downsampled to 250 Hz. Further processing was performed using MNE-Python (Gramfort et al., 2013) and FieldTrip (Oostenveld et al., 2011). For each participant, data were concatenated across the 5 recording blocks. To reduce the influence of eye movement and cardiac activity related artefacts, an independent component analysis (ICA, FastICA method Hyvarinen, 1999) was performed. Before ICA fitting, the data were filtered between 1-45 Hz, whitened (decorrelated and scaled to unit variance), and their dimensionality reduced by means of principal component analysis (PCA). The first n PCA components explaining the cumulative variance of 0.9 of the data were entered in the ICA decomposition. The computed ICA filters were applied on the concatenated raw data. Between-participant differences in head position were compensated for by transforming MEG data from each participant to the mean sensor array across participants using MaxFilter 2.2. Finally, epochs were extracted time locked to the onset of the speaker’s mouth movement (0-5 s).

To measure neural phase coherence with auditory and visual speech, we created two speech signals from each of the stimulus video clips: (i) the acoustic envelope of the auditory speech and (ii) the time course of the instantaneous area of the speakers’ lip aperture (Fig. 1B). For each sentence, the amplitude envelope of the auditory speech signal was extracted using custom Matlab code following a standard sequence of steps similar to noise vocoding: full wave rectification and low-pass filtering at 30 Hz. The lip aperture envelope of the speaker was computed using custom Matlab code from (Park et al., 2016). Briefly, for each frame, the lip contour of the speaker was extracted and the area within the lip contour was calculated. To match the amplitude and sampling rate of the auditory and visual speech envelopes to the MEG signal, they were scaled by a factor of 10^-10^ and 10^-15^ respectively and resampled to 250 Hz.

For each participant, high-resolution structural MRI images (T1-weighted) were obtained using a GRAPPA 3-D MPRAGE sequence (resolution time = 2250 msec, echo time = 2.99 msec, flip angle = 9°, and acceleration factor = 2) on a 3T Tim Trio MR scanner (Siemens) with 1 × 1 × 1 mm isotropic voxels. MRI images were segmented with FreeSurfer (Fischl, 2012) using the default parameter settings.

### Speech signal analysis and statistics

To relate the auditory envelope (i.e., the broadband acoustic speech envelope) to the lip aperture envelope (i.e., the visual component of speech), we computed their coherence. First, the data were transformed to frequency domain using a fast Fourier transform (FFT) applied to the entire auditory and visual speech signals using a Hanning window, producing spectra with a frequency resolution of 0.5 Hz between 0.5 and 20 Hz. Then, the cross-spectral density was computed between the two signals. Finally, the coherence was computed between the two signals 𝑖 and 𝑗 for each frequency 𝑓 as the magnitude of the cross-spectral density (𝐶𝑆𝐷) divided by the square root of the power spectra of both signals:

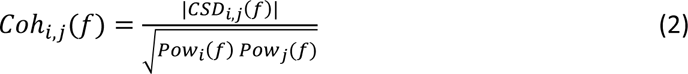

We fitted a linear function to the log-log transformed power spectra of the auditory and visual speech signals to illustrate their natural 1/𝑓 noise profile (Fig. 2A, Chandrasekaran et al., 2009).

**Figure 2.**
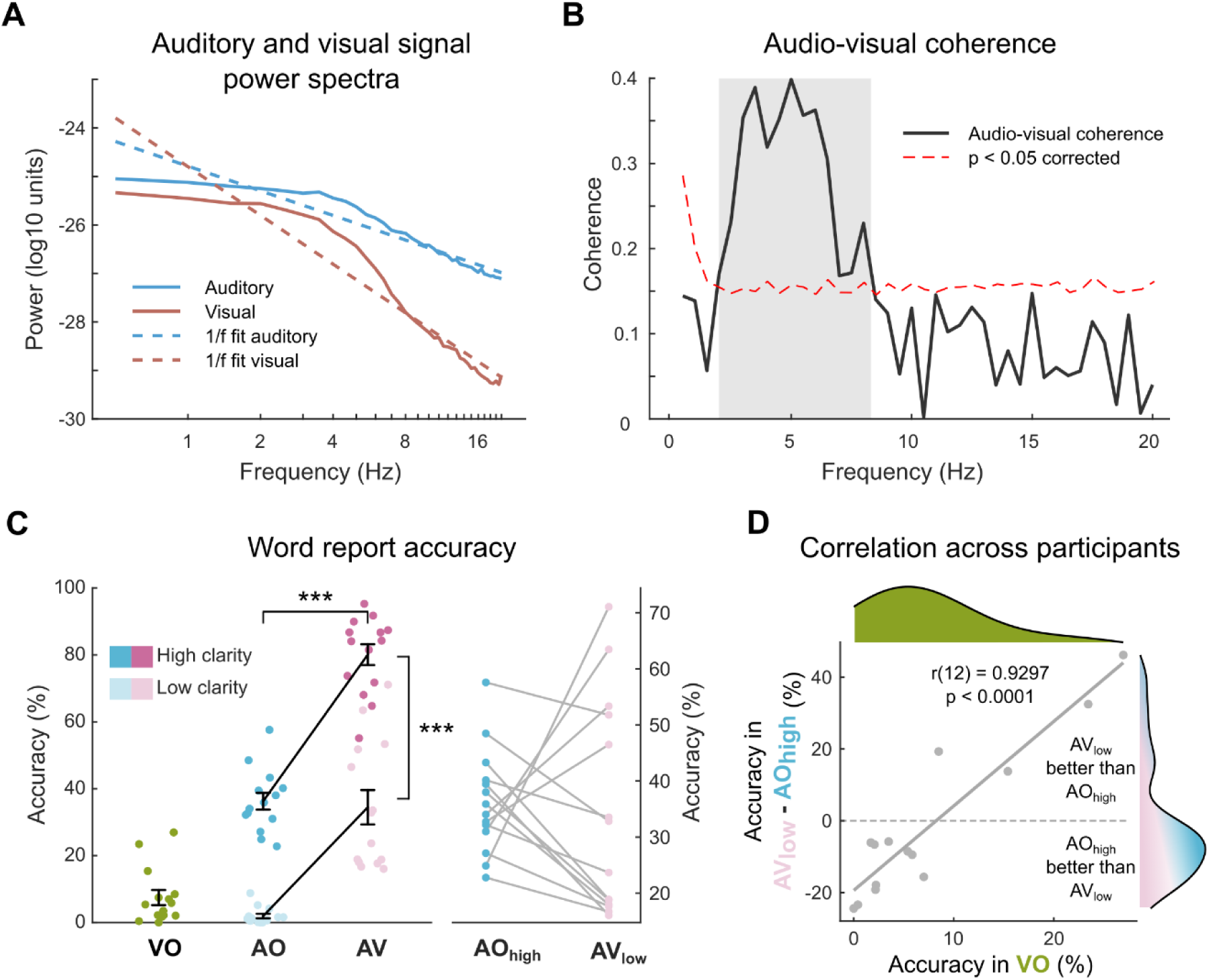
Stimulus characteristics and behavioural results. **A.** Power spectra and fitted 1/f noise profiles of auditory and visual speech signals. **B.** Coherence between auditory and visual speech signals across frequencies. Dashed lines indicate significance level, corrected for multiple comparisons, shading indicates 2-8 Hz range with greater-than-chance audio-visual coherence. **C.** Group level word report accuracies (mean ± SEM) overlayed with individual data across conditions. The right side of the figure shows the individual differences between AO_high_ and AV_low_ conditions matched for overall intelligibility. **D.** Correlation across participants between the measure of audio-visual benefit (word report difference between AV_low_ and AO_high_) and lip reading ability (word report accuracy in VO). Marginal distributions for the two variables are displayed at the top and right hand side, respectively. (AO: audio-only, AV: audio-visual, VO: visual-only, *** p < 0.001)

We performed permutation tests to establish at which frequencies the auditory and visual speech signals were coherent. We randomly permuted the assignment of auditory and visual speech signals 5000 times generating a null distribution of permuted coherence values. Then, for each frequency, we computed the proportion of permuted coherence values greater than the observed coherence value (Fig. 2B) equivalent to a one-tailed p-value. To control for multiple comparisons across frequencies, we applied Bonferroni correction.

### Behavioural data analysis and statistics

Word report accuracies for each participant were entered in a 2 (acoustic clarity high vs. low) x 2 (visual speech present vs. absent) repeated measures ANOVA and the main effects of acoustic clarity and availability of visual speech were reported (Fig. 2C). We also computed the Pearson correlation between word report accuracy in VO condition (as an index of lip reading ability) and the word report accuracy difference between AV_low_ and AO_high_ conditions (expressing the relative benefit that participants received from providing ‘visual’ or ‘auditory’ speech signals compared to the most difficult AO_low_ condition). Importantly these two methods are independent under the null hypothesis and show considerable variation across participants (Fig. 2D).

To test the distribution of the measure of audio-visual benefit for unimodality, we computed Hartigan’s dip statistic (Hartigan and Hartigan, 1985) and its corresponding p-value. The dip statistic expresses the maximum distance between the empirical distribution and the best fitting unimodal distribution on a scale between 0 and 1. The null hypothesis of the test is unimodal distribution; hence a significant test statistic is interpreted as evidence for non-unimodal, i.e., at least bimodal distribution.

### MEG data analysis and statistics

To measure neural phase coherence with auditory and visual speech at the sensor level, we computed the coherence between the magnetometers and the auditory and visuals speech signals, respectively, in line with previous studies (Peelle et al., 2013). This analysis was conducted to establish MEG-auditory (i.e., between neural and auditory envelope) and MEG-visual coherence effects (i.e., coherence between the neural signal and visual envelope). Basic MEG-auditory and MEG-visual coherence was established in analyses including all conditions with an auditory (i.e., AO_high_, AO_low_, AV_high_, AV_low_) or visual signal (i.e., AV_high_, AV_low_, VO), respectively. Coherence was computed between 1-20 Hz at 1 Hz increments. For each participant, we also computed the coherence for 100 random pairings of auditory/visual and neural data, making sure that none of the auditory/visual signals were paired with their original neural signal pair. These permutations were then averaged for each participant to produce coherence maps which provide an estimate of the baseline coherence values that can be expected by chance. The difference between this permuted coherence measure and observed coherence for the true pairing of auditory/visual signals and neural responses is normally distributed, no longer bounded between 0 and 1 and is expected to be zero under the null hypothesis of no coherence between sensory and neural signals. This is a more suitable dependent variable for statistical tests and hence we report the difference between true and permuted coherence in all analyses. We performed cluster-based non-parametric permutation tests (Maris and Oostenveld, 2007) to test if there is reliable coherence to auditory and/or visual speech above chance (i.e., true - permuted coherence > 0). The frequency range of the test was restricted to frequencies exhibiting greater-than-chance coherence between the auditory and visual envelopes, (i.e., 2-8 Hz, see Fig. 2B and results).

Source space analysis was performed in MNE-Python (Gramfort et al., 2013). For each participant, MEG sensor positions were co-registered with the individual MRI images and visually verified. The forward solution was computed using a one-layer boundary element model (BEM) based on each participant’s inner skull mesh obtained from the FreeSurfer segmentation of individual anatomical MRI images. We used dynamic imaging of coherent sources (DICS, Gross et al., 2001) to determine the spatial distribution of brain areas coherent to the auditory and visual speech signals (Peelle et al., 2013). Cortico-auditory and cortico-visual coherence source maps were computed at 4096 vertices in each hemisphere, in increments of 0.2 Hz between 2-6 Hz and averaged across frequencies before group statistics. The frequency range was based on the frequency extent of the significant clusters observed in the basic MEG-auditory and MEG-visual coherence effects in sensor space. It has been shown previously that the auditory and visual speech signals are coherent in audio-visual speech (Chandrasekaran et al., 2009), a finding which we also confirmed for our stimulus materials. It is thus possible that some of the observed cortico-auditory coherence effects could be accounted for by the visual signal (e.g., in occipital cortex), and vice-versa for cortico-visual coherence in auditory cortex. To rule out this possibility, we computed partial coherence (Rosenberg et al., 1998) between neural and auditory/visual speech signals (i.e., cortico-auditory coherence after removing coherence explained by the visual signal and cortico-visual coherence after removing coherence explained by the auditory signal) which we once more compared to null distributions based on computing partial coherence for 100 random combinations of auditory/visual and neural data.

To rule out the possibility that our results were biased by imbalanced number of trials between the conditions being compared (e.g., when comparing AV and VO conditions, the latter only included 55 trials, half of the former condition), we re-computed the coherence/partial coherence for the more abundant condition with subsampling. More specifically, we randomly sampled trials from the more abundant condition without replacement to match the number of trials in the less abundant condition and computed the coherence/partial coherence. This procedure was repeated 100 times with a new random sample of trials, then the resulting coherence or partial coherence values were averaged across the 100 repetitions.

We defined two regions of interest (ROI) based on the anatomical parcellation by (Destrieux et al., 2010): superior temporal gyrus (STG, i.e., lateral aspect of the superior temporal gyrus) and occipital cortex (OCC, including inferior occipital gyrus and sulcus, cuneus, middle and superior occipital gyri, occipital pole, middle and superior occipital sulci, lunatus sulcus, and transverse occipital sulcus). For each participant, true and permuted coherence values were averaged across vertices within each ROI and were entered in group level statistics.

All statistical tests on MEG data were performed at the second, between-subjects level. In the sensor space analysis we used cluster-based non-parametric permutation tests (Maris and Oostenveld, 2007) based on the t-statistic to identify clusters in the 102 channels x 7 frequencies (2-8 Hz) data space that exhibit greater true than permuted coherence between neural and auditory/visual speech signals. These second level tests were based on 5000 permutations, using a cluster-defining threshold of p < 0.05 and a test threshold of p < 0.05.

For statistical analysis in source space the individual source coherence maps were first morphed onto the average brain provided by FreeSurfer (‘fsaverage’). In the whole-brain source analysis we performed the same cluster-based permutation tests. We used a more stringent cluster defining threshold of p < 0.005 for assessing overall cortico-auditory and cortico-visual coherence; this assisted with the clarity of visualisation given the very reliable results shown for this analysis. We used a cluster defining threshold of p < 0.05 with cortico-auditory and cortico-visual coherence after partialling out visual and auditory signals, respectively.

For the ROI analyses we first subtracted the permuted whole-brain partial coherence from the true partial coherence maps in each condition. Then we averaged the vertex-level true-minus-permuted coherence values within each bilateral ROI and performed paired t-tests across participants. We computed a one-tailed p-value when comparing true-permuted partial coherence to 0 as true coherence is expected to be greater than permuted. For comparisons of true-permuted partial coherence between two conditions, we computed a two-tailed p-value.

### Code and data availability

All analysis code and data will be available for download upon the publication of the manuscript.

## Results

### Audio-visual properties of sentences

First, we characterized the properties of the speech signals in the set of sentence stimuli. We performed a frequency analysis on the auditory and visual speech signals and fitted a 1/f function to the average power spectra across sentences to indicate the expected noise profile (Voss and Clarke, 1975) (Fig. 2A). The residual power spectra showed a maximum between 2-8 Hz consistent with previous findings (Chandrasekaran et al., 2009; Peelle et al., 2013). Previous results also indicated that the auditory and visual speech signals are closely coupled (Chandrasekaran et al., 2009), hence we conducted a coherence analysis between the auditory and visual speech signals (Fig. 2B). The coherence spectrum also shows a clear peak between 2-8 Hz (see shaded area in Fig. 2B). Permutation tests confirmed this observation, concluding that coherence between auditory and visual speech signals was above chance between 2-8 Hz (p < 0.05, corrected for multiple comparisons). This analysis as well as previous literature informed our subsequent analyses of coherence between neural and speech signals, allowing us to narrow our focus to the frequency range for which audio-visual interactions are expected.

### Behavioural results

We collected and scored participants’ word reports for accuracy and computed the average word report accuracies across sentences separately for each condition (Fig. 2C). As expected, word report accuracy was lowest in the AO_low_ condition (mean ± SEM: 1.95% ± 0.67%; similar to 1 channel vocoded speech, see Peelle et al., 2013; Sohoglu et al., 2014; Zoefel et al., 2018). Both increased acoustic clarity and visual speech improved word report accuracy as indicated by significant main effects (acoustic clarity: low (mean ± SEM): 18.20% ± 5.70% vs. high (mean ± SEM): 58.16% ± 6.57%, F(1,13) = 221.163, p < 0.0001; visual speech: AO (mean ± SEM): 19.10% ± 5.00% vs. AV (mean ± SEM): 57.25% ± 7.47%, F(1,13) = 93.969, p < 0.0001). Their interaction was also significant (F(1,13) = 10.427, p = 0.0066), but this might reflect non-linearities in the accuracy measure which in some conditions approached maximum and minimum values. Simple effects of acoustic clarity for AO and AV speech, and simple effects of visual speech for high and low clarity speech were all reliable (t(13) > 6.025, p < 0.0001). Word report accuracy in visual only speech (VO) was also relatively low but showed higher variability than AO_low_ (mean ± SEM: 7.47% ± 2.28%).

Importantly, our manipulation of acoustic clarity based on data from a pilot experiment ensured that sentences in AO_high_ and AV_low_ conditions were approximately equally intelligible (AO_high_ (mean ± SEM): 36.26% ± 2.50%; AV_low_ (mean ± SEM): 34.45% ± 5.13%). Bayesian paired t-test on word report accuracies between these two conditions provided approximately 3.5 times stronger evidence in favour of the null-hypothesis of no difference than the alternative (t(13) = -0.314, p = 0.7587, BF01 = 3.55). These conditions therefore enabled us to compare two conditions with different sources of additional speech information (i.e., increased acoustic clarity in AO_high_ and low acoustic clarity supported by visual speech in AV_low_) with minimal intelligibility confound (Fig. 2C and D). This analysis revealed marked differences across participants: some benefitted more from the additional visual speech signal than from the increased acoustic clarity (compare lines with positive and negative slopes in Fig. 2C right). We quantified this by computing the difference in word report accuracies between AV_low_ and AO_high_. This measure of ‘audio-visual benefit’ appeared to follow a bimodal distribution (see marginal distributions in Fig. 2D). To confirm our visual observation, we tested the data for unimodality using Hartigan’s dip statistic (Hartigan and Hartigan, 1985), however the results (D = 0.0736, p = 0.7649) did not provide reliable evidence for non-unimodal distribution perhaps because of our relatively small sample of listeners. Nevertheless, the measure of audio-visual benefit correlated with participant’s lipreading ability (as indexed by word report accuracy in VO). The better participants were able to lipread (i.e., report words in silent speech), the more they benefitted from the additional visual speech signal (Pearson’s r(12) = 0.9297, p < 0.0001, Fig. 2D).

### Coherence between neural signals and speech

First, we established the basic coherence between MEG and the auditory/visual speech signals in sensor space. We computed the coherence between signals recorded from each magnetometer and the auditory speech signal in all conditions containing auditory speech signals (i.e., AO_high_, AO_low_, AV_high_ and AV_low_), and the coherence between each magnetometer and the visual speech signal in all conditions containing visual speech signals (i.e., AV_high_ and AV_low_, VO). We also computed the corresponding ‘permuted’ coherence maps by randomly pairing magnetometer signals with auditory/visual speech signals from other trials (see MEG data analysis for details). This estimated a null distribution for the degree of coherence that can be expected by chance. We contrasted the true and permuted coherence values using a cluster-based permutation test across participants (Maris and Oostenveld, 2007). This analysis revealed that brain responses phase locked to both auditory and visual speech as indicated by significantly above-chance coherence between magnetometer and auditory/visual speech signals (Fig. 3A). The cluster-based permutation tests revealed bilateral clusters phase locked to auditory speech (p = 0.0002, corrected) and a posterior cluster phase locked to visual speech (p = 0.0002, corrected), both spanning frequencies between 2-6 Hz (Fig. 3A). Based on these results, we further narrowed our frequency range of interest to 2-6 Hz in subsequent analyses of neural sources.

**Figure 3.**
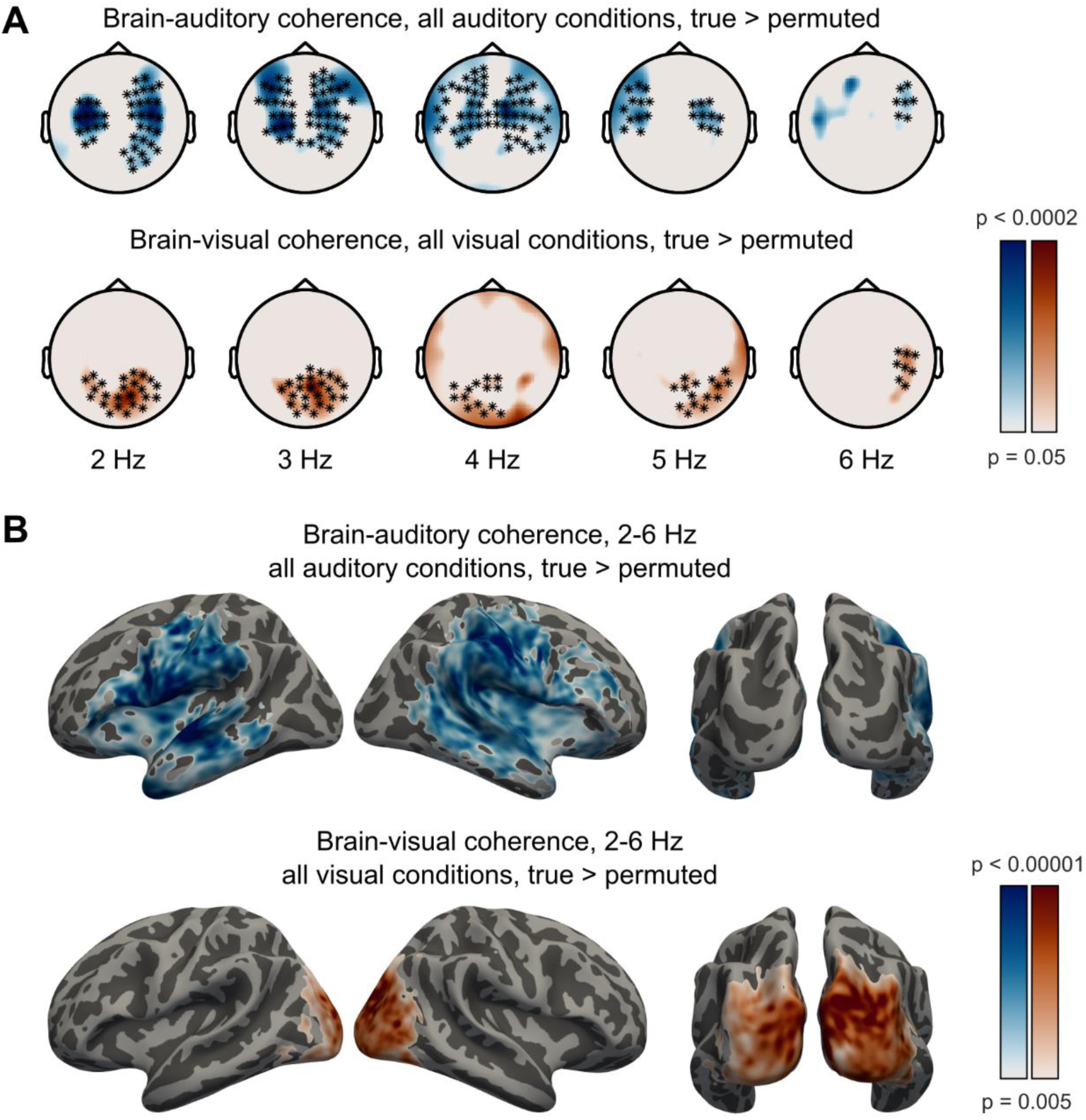
Coherence between neural and speech signals. **A.** Sensor topographies show MEG-auditory coherence above permutation-null baseline in all auditory conditions (top) and MEG-visual coherence above baseline in all visual conditions (bottom) across frequencies. Markers show clusters which were statistically significant (p = 0.0002) in the cluster-based permutation test (one continuous cluster each, spanning 2-6 Hz for both MEG-auditory and MEG-visual coherence). **B.** Source maps show cortico- auditory coherence above permutation-null baseline in all auditory conditions (top) and cortico-visual coherence above baseline in all visual conditions (bottom) averaged across frequencies between 2-6 Hz. Effects shown are whole-brain cluster corrected (p < 0.05) based on cortical sources exceeding a vertex-level threshold of p<0.005 (inset colour scale).

To reveal which brain areas phase locked to auditory and visual speech signals, we conducted a whole-brain analysis on source localized MEG responses (Fig. 3B). We performed cluster-based permutation tests on source coherence maps averaged between 2-6 Hz to compare true and permuted coherence between neural and auditory/visual speech signals across participants. We observed greater-than-chance phase coherence with both auditory and visual speech signals as indicated by significant, non-overlapping clusters centred on auditory and visual cortex (p < 0.05, whole-brain corrected). Bilateral temporal, parietal, and inferior-frontal areas phase locked to auditory speech, whereas bilateral occipital areas phase locked to visual speech.

Next, we investigated which brain areas phase locked more strongly to the auditory than the visual speech signal and vice versa. In this analysis we used only the sentences containing both auditory and visual speech signals (i.e., the AV_high_, and AV_low_ conditions). Importantly, the auditory and visual speech signals are coherent with each other (Chandrasekaran et al., 2009) (also see Fig. 2B). Hence, to rule out the possibility that the observed neural coherence with one speech signal (e.g., auditory) is explained by the other speech signal (e.g., visual), we computed the partial coherence (Rosenberg et al., 1998), i.e., the neural coherence with the auditory (respectively visual) speech signal after removing influences of the visual signal and vice-versa (see Park et al., 2016, for a similar approach). Fig. 4A shows whole-brain maps of auditory > visual and visual > auditory partial coherence effects in AV conditions. Temporal, frontal, and parietal areas were phase locked more strongly to auditory than visual speech, whereas occipital areas were phase locked more strongly to visual than auditory speech, as shown by cluster-based permutation tests (p < 0.05, whole-brain corrected).

**Figure 4.**
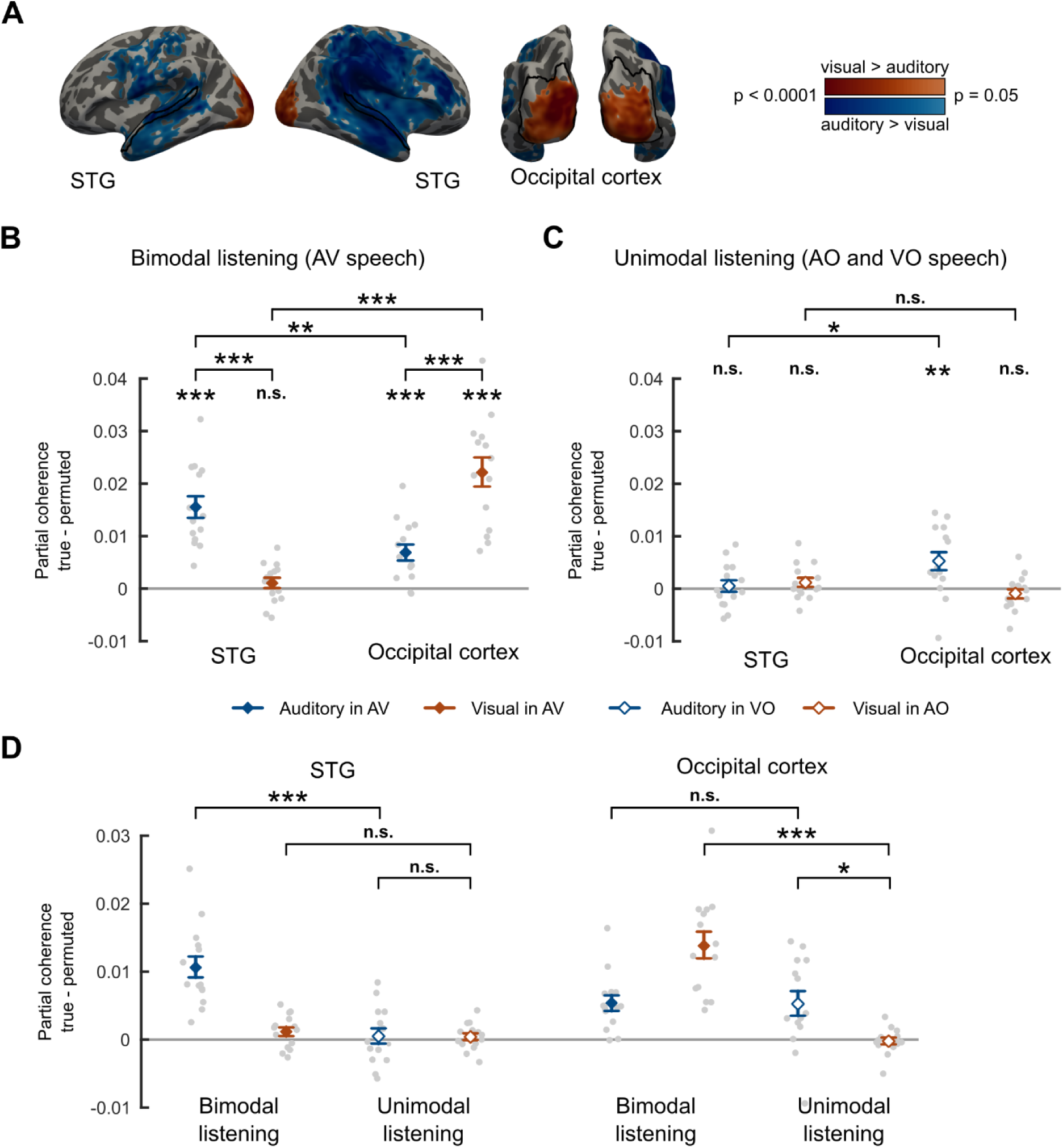
Partial auditory and visual coherence in low and high clarity audio-visual conditions (AV_low_, AV_high_). **A.** Whole-brain maps of partial coherence contrasts averaged between 2-6 Hz: auditory > visual and visual > auditory in bimodal listening conditions (i.e., audio-visual (AV) speech). Effects shown are whole-brain corrected (p < 0.05) using a vertex level threshold of p<0.05 (see colour scale). Black outlines mark the regions used in the ROI analysis. **B.** Group level auditory and visual partial coherence showing effects of modality in bimodal listening conditions. Coherence computed with respect to the permutation null baseline in STG and Occipital cortex. **C.** Group level partial coherence showing cross-modal effects in unimodal listening conditions (i.e., auditory partial coherence in silent speech (VO) and visual partial coherence in audio-only speech (AO)). Only within-condition comparisons are permitted for this data due to the different number of trials in AO and VO conditions. D. Group level partial coherence with respect to the permutation null baseline across ROIs in unimodal and bimodal listening conditions with AO and AV conditions resampled to match the number of trials in VO (necessary for between-condition comparisons). Diamond markers and error bars represent mean ± SEM over participants, suitable for comparisons with 0, but not for repeated-measures comparisons between conditions. Grey dots represent individual data. (ROI: region of interest, STG: superior temporal gyrus, n.s.: not significant, * p < 0.05, ** p < 0.01, *** p < 0.001)

Next, we examined auditory and visual phase coherence in auditory and visual cortical areas involved in audio-visual speech processing. We defined two regions of interest (ROI) based on the anatomical parcellation by Destrieux et al., 2010. For classical visual areas, we defined an occipital cortex ROI covering bilateral occipital areas (for details see MEG data analysis, also Fig. 4A bottom). To define a speech-responsive auditory ROI we used the bilateral superior temporal gyrus (STG, Fig. 4A top). We computed the difference between true and permuted cortico-auditory and cortico-visual partial coherence at each vertex and averaged them across vertices within each ROI. Fig. 4B shows the individual partial auditory and visual coherence values in audio-visual conditions with respect to the permutation null baseline for these ROIs. As expected, in STG, we observed significant phase coherence with the auditory speech signal (mean ± SEM = 0.0155 ± 0.0021, t(13) = 7.397, one-tailed p < 0.0001), and that STG phase locked to auditory speech significantly more strongly than visual speech (t(13) = 6.448, two-tailed p < 0.0001). However, we did not find evidence of significant phase coherence between STG and visual speech once auditory signals were partialled out (mean ± SEM = 0.0010 ± 0.0010, t(13) = 1.022, one-tailed p = 0.1627). In occipital cortex, we observed above chance phase coherence with visual signals as indicated by a significant one-sample t-test of true - permuted partial visual coherence against 0 (mean ± SEM = 0.0221 ± 0.0028, t(13) = 7.801, one-tailed p < 0.0001). Furthermore, occipital cortex showed stronger phase coherence with visual than auditory speech as indicated by a paired t-test between auditory and visual partial coherence (t(13) = 4.469, two-tailed p = 0.0006). This finding is in line with the overlap shown in Fig. 4A between our ROIs, and the clusters of modality-specific partial coherence shown in whole-brain analyses. However, surprisingly, we also observed that occipital cortex phase locked to auditory speech, even when visual coherence was partialled out (mean ± SEM = 0.0069 ± 0.0016, t(13) = 4.379, one-tailed p = 0.0004). A repeated measures ANOVA with factors coherence modality (auditory vs. visual) and ROI (STG vs. occipital cortex) revealed a significant interaction between the two factors (F(1,13) = 52.289, p < 0.0001). Pairwise comparisons between ROIs confirmed that phase-locking to auditory speech was stronger in STG (t(13) = 3.837, two-tailed p = 0.0021) and phase-locking to visual speech was stronger in occipital cortex (t(13) = 7.583, two-tailed p < 0.0001). These results indicate that in bimodal listening conditions the auditory and visual envelopes of speech are tracked strongest in their modality-preferred cortices (i.e., auditory in STG and visual in occipital cortex).

We also assessed whether phase-locking to the auditory speech envelope is observed during silent lip reading (VO condition) and conversely, whether phase-locking to the visual speech envelope is observed during audio-only speech (AO conditions) in our ROIs. Hence, we computed partial auditory coherence in VO and partial visual coherence in AO conditions with respect to the permutation baseline in STG and occipital cortex (Fig. 4C). In occipital cortex, we found evidence of phase-locking to auditory speech signals during responses to silent visual speech (mean ± SEM = 0.0053 ± 0.0018, t(13) = 2.905, one-tailed p = 0.0061) but no evidence of responses to visual speech signals in response to audio-only speech (mean ± SEM = -0.0009 ± 0.0009, t(13) = -1.011, one-tailed p = 0.1652). In STG, we did not find reliable evidence of phase tracking of auditory speech in VO, nor visual speech in AO (t(13) < 1.293, one-tailed p > 0.1093). Pairwise comparisons between ROIs revealed that phase-locking to the auditory speech envelope during silent lip reading was stronger in occipital cortex than in the STG (t(13) = 2.223, two-tailed p = 0.0445) but we found no difference between these ROIs in phase-locking to the visual speech envelope during audio-only speech (t(13) = 1.930, two-tailed p = 0.0757). These results suggest differences between the tracking of auditory and visual speech in unimodal and bimodal listening conditions that we now explore further.

To formally compare speech tracking between uni- and bimodal listening conditions (Fig. 4D), we sub-sampled the AV and AO conditions to ensure that these conditions include the same number of trials as the VO condition. We then performed a repeated measures ANOVA with 3 factors: listening condition (unimodal vs. bimodal), coherence modality (auditory vs. visual) and ROI (STG vs. occipital cortex) on this sub-sampled data. This showed a significant 3-way interaction (F(1,13) = 81.871, p < 0.0001) indicating differences between auditory and visual cortex ROIs in phase-locking to auditory and visual speech signals during unimodal and bimodal listening. To characterise this interaction we conducted separate repeated measures ANOVAs in each ROI with factors listening condition and coherence modality that confirmed significant two-way interactions of listening condition and coherence modality in both ROIs (STG: F(1,13) = 52.096, p < 0.0001; occipital cortex: F(1,13) = 21.137, p = 0.0005) though as shown by the three-way interaction, these differ between ROIs. Pairwise comparisons revealed that in STG auditory phase-locking is stronger in bimodal than unimodal listening (t(13) = 8.063, two-tailed p < 0.0001). The same difference between bimodal and unimodal listening is observed for visual phase-locking in occipital cortex (t(13) = 7.400, two-tailed p < 0.0001).

The above three-way interaction indicates that the STG and occipital cortex differ in their response to speech envelopes in their non-preferred modality. Specifically, in the STG there is no phase-locking to visual speech envelopes during processing of unimodal or bimodal auditory speech signals (comparisons with null distributions reported above and in Fig. 4B and C) and no reliable difference between phase-locking to visual speech in bimodal and unimodal conditions (visual in AV vs. visual in AO difference t(13) = 0.857, two-tailed p = 0.4072) or between responses to auditory and visual information in unimodal conditions (auditory in VO vs. visual in AO, t(13) = 0.078, two-tailed p = 0.9388). Thus, the STG does not phase lock to visual speech signals in either AV or AO conditions, and only phase locks to auditory signals when these are physically present. Conversely, occipital cortex shows reliable phase locking to auditory speech envelopes irrespective of whether these envelopes are present (AV condition) or absent (VO condition, see stats in Fig. 4B and C). Furthermore, there is no reliable difference in auditory phase-locking between bimodal (AV) and unimodal (VO) listening conditions (t(13) = 0.080, two-tailed p = 0.9372) and phase-locking to (absent) auditory speech signals in VO conditions is greater than that seen for (absent) visual speech signals in AO conditions (t(13) = 2.618, two-tailed p = 0.0213). Hence, occipital cortex shows a greater degree of phase-locking to speech signals in the non-preferred modality than the STG; this cross-modal influence is observed during listening conditions that involve audio-visual benefit (AV) or lip-reading (VO).

Next, we compared phase-locking to auditory speech signals during bimodal (AV) and unimodal (AO) listening in visual and auditory ROIs. As previously shown in Fig. 4B, above chance auditory phase locking is observed in occipital cortex when both auditory and visual signals are present (i.e., AV conditions) even when statistical influences of correlated visual signals are excluded (using partial coherence analysis). Comparison of AV and AO conditions further confirms that auditory phase-locking in occipital cortex is enhanced by the presence of visual speech: partial coherence with auditory signals was greater in AV conditions than in AO conditions (t(13) = 3.017, two-tailed p = 0.0099, see Fig. 5A, B). Whole-brain comparisons of these two conditions are reported subsequently (see Fig. 5A, Table 1). To ensure that additional auditory phase locking in occipital cortex was due to the presence of visual speech and not accompanying differences in intelligibility we also compared partial coherence in AV_low_ and AO_high_ conditions that are matched for average intelligibility (see Fig. 2C). This comparison confirmed stronger phase-locking to auditory speech signals in occipital cortex when visual speech is present (t(13) = 2.552, two-tailed p = 0.0241, see Fig. 5D). However, this effect did not correlate with individual differences in intelligibility (i.e., our measure of audio-visual benefit, Fig. 2D) between these otherwise matched conditions (Pearson’s r(12) = -0.272, two-tailed p = 0.3476). Despite this beneficial effect of visual speech, auditory phase-locking remains reliable in occipital cortex even when only auditory speech signals are present. We see significantly greater partial auditory coherence in AO conditions (i.e., AO_high_ and AO_low_ combined) than in a null baseline computed by permuting the assignment of speech envelopes to MEG data (mean ± SEM = 0.0026 ± 0.0009, t(13) = 2.906, one-tailed p = 0.0061, see Fig. 5B).

**Figure 5.**
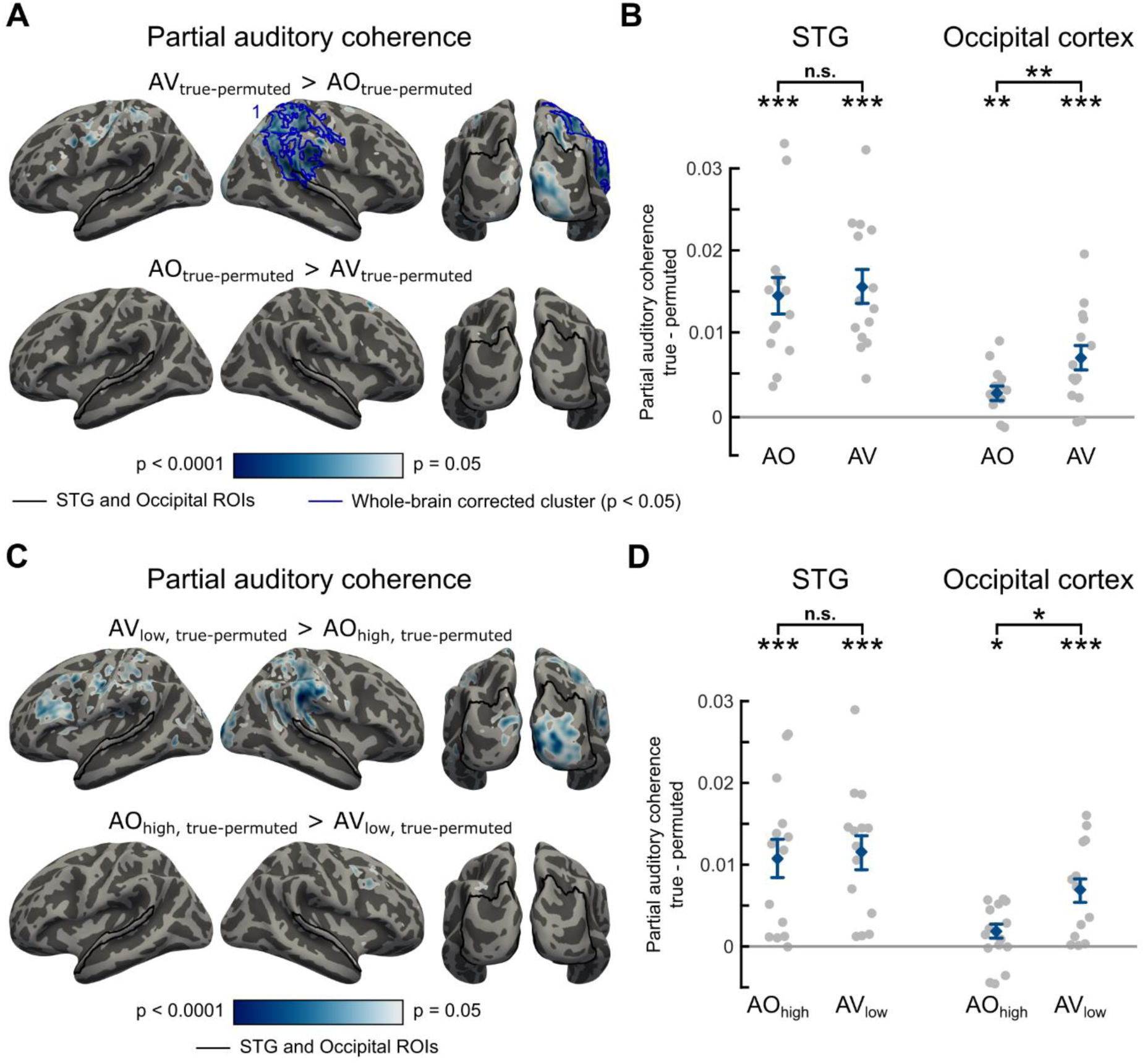
Cross-modal influences on auditory phase locking in auditory and visual areas. Effect of visual speech availability on auditory partial coherence, whole-brain (A, C) and ROI-based analysis (B, D). **A.** Whole-brain maps show significant effects of visual speech availability on partial auditory coherence with respect to permutation-derived null baseline averaged between 2-6 Hz (uncorrected p-values). Black outlines indicate STG and occipital ROIs, blue outlines indicate whole-brain corrected clusters (p < 0.05). **B.** Graphs depict the mean auditory partial coherence with respect to the permutation null baseline averaged between 2-6 Hz in AV and AO conditions across ROIs. Diamonds and error bars represent mean ± SEM, grey dots represent individual data points. **C.** Whole-brain maps show significant effects of visual speech availability (controlling for speech intelligibility) on auditory partial coherence with respect to permutation-derived null baseline averaged between 2-6 Hz (uncorrected p-values). Black outlines indicate STG and occipital ROIs, red outlines indicate whole-brain corrected clusters (p < 0.05). **D.** Graphs depict the mean auditory partial coherence with respect to the permutation null baseline averaged between 2-6 Hz in AV_low_ and AO_high_ conditions across ROIs. Diamonds and error bars represent mean ± SEM, grey dots represent individual data points. The clusters supporting the whole-brain corrected significant results are numbered and further details are presented in Table 1. (ROI: region of interest, STG: superior temporal gyrus, AO: audio-only, AV: audio-visual, n.s.: not significant, * p < 0.05, ** p < 0.01, *** p < 0.001)

**Table 1.**
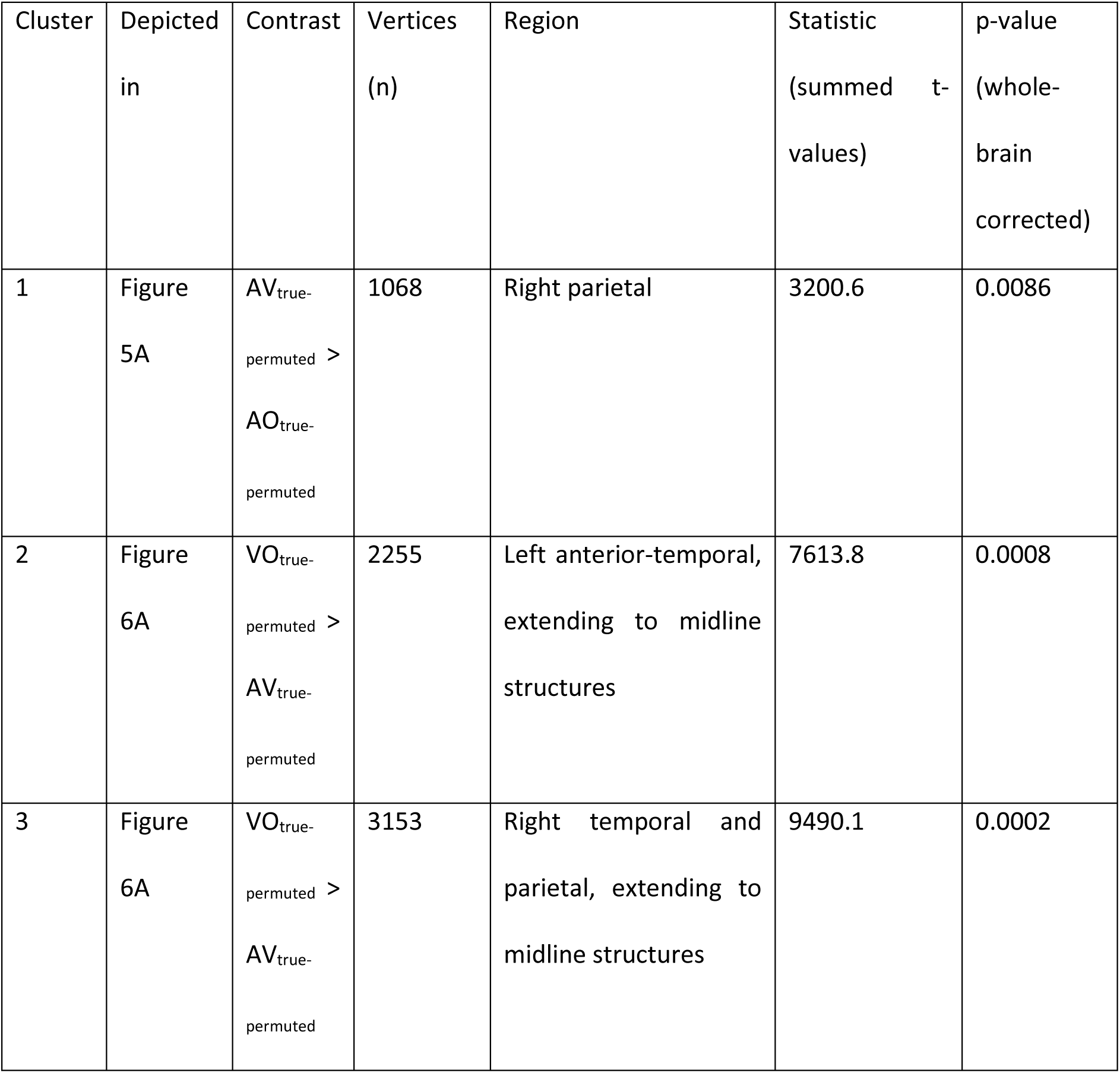
Descriptions and summary statistics of whole-brain analysis clusters.

These cross-modal influences were again numerically less apparent in our auditory ROI. Comparison of AV and AO conditions showed no evidence that auditory partial coherence in the STG was influenced by the presence of visual speech when high and low clarity speech conditions were combined (t(13) = 0.809, two-tailed p = 0.4329, see Fig. 5B) or when comparing intelligibility-matched AV_low_ and AO_high_ conditions (t(13) = 0.370, two-tailed p = 0.7173, Fig. 5D). Partial auditory coherence in the STG is greater than the permutation baseline both in AV conditions (shown previously in Fig. 4B) and in AO conditions (mean ± SEM = 0.0145 ± 0.0023, t(13) = 6.258, one-tailed p < 0.0001, Fig. 5B). However, a two-way ANOVA testing for the interaction between stimulus modality (AV vs. AO) and ROI (STG vs. Occipital cortex) failed to show a significant interaction either for all AV and AO trials (F(1,13) = 2.954, p = 0.1094), or for intelligibility matched AVlow and AOhigh conditions (F(1,13) = 2.693, p = 0.1248). Hence, we have only limited evidence that visual enhancement of auditory phase-locking is specific to occipital regions.

Bimodal and unimodal listening conditions can also be assessed by comparing phase-locking to visual speech in AV conditions (as before) with VO conditions in which auditory signals are absent. For these comparisons we sub-sampled trials from the two AV conditions to ensure that the number of trials matched the VO condition. Significant phase locking to visual speech signals in STG was seen in the presence of both auditory and visual speech that narrowly exceeded the permutation baseline (mean ± SEM = 0.0012 ± 0.0007, t(13) = 1.783, one-tailed p = 0.049, see Fig. 6B). We had previously observed that this effect was not reliable when the full set of AV trials was analysed (see Fig. 4B) and hence interpret this finding with caution. Nonetheless, comparisons of results from the subsampled AV condition and VO condition are informative since this comparison is matched for the number of trials. We observed reliable visual phase coherence in the STG in the absence of auditory speech (i.e., in the VO condition). This is indicated by significant partial visual coherence with respect to the permutation null baseline (mean ± SEM = 0.0056 ± 0.0013, t(13) = 4.499, one-tailed p = 0.0003). Indeed, visual phase locking in STG was greater in the absence of auditory speech than when it was present (VO > AV, (t(13) = 4.185, two-tailed p = 0.0011, Fig. 6B). This was the opposite outcome to that previously reported for auditory phase locking in occipital cortex (Fig. 5D). Greater phase-locking to visual speech signals in auditory cortex during silent lip-reading might suggest that speech signals are reinstated or filled-in from visual input. We will expand on this finding in the discussion. However, this effect did not reliably correlate with individual differences in lip reading (i.e. word report in VO conditions, Pearson’s r(12) = -0.220, two-tailed p = 0.450).

**Figure 6.**
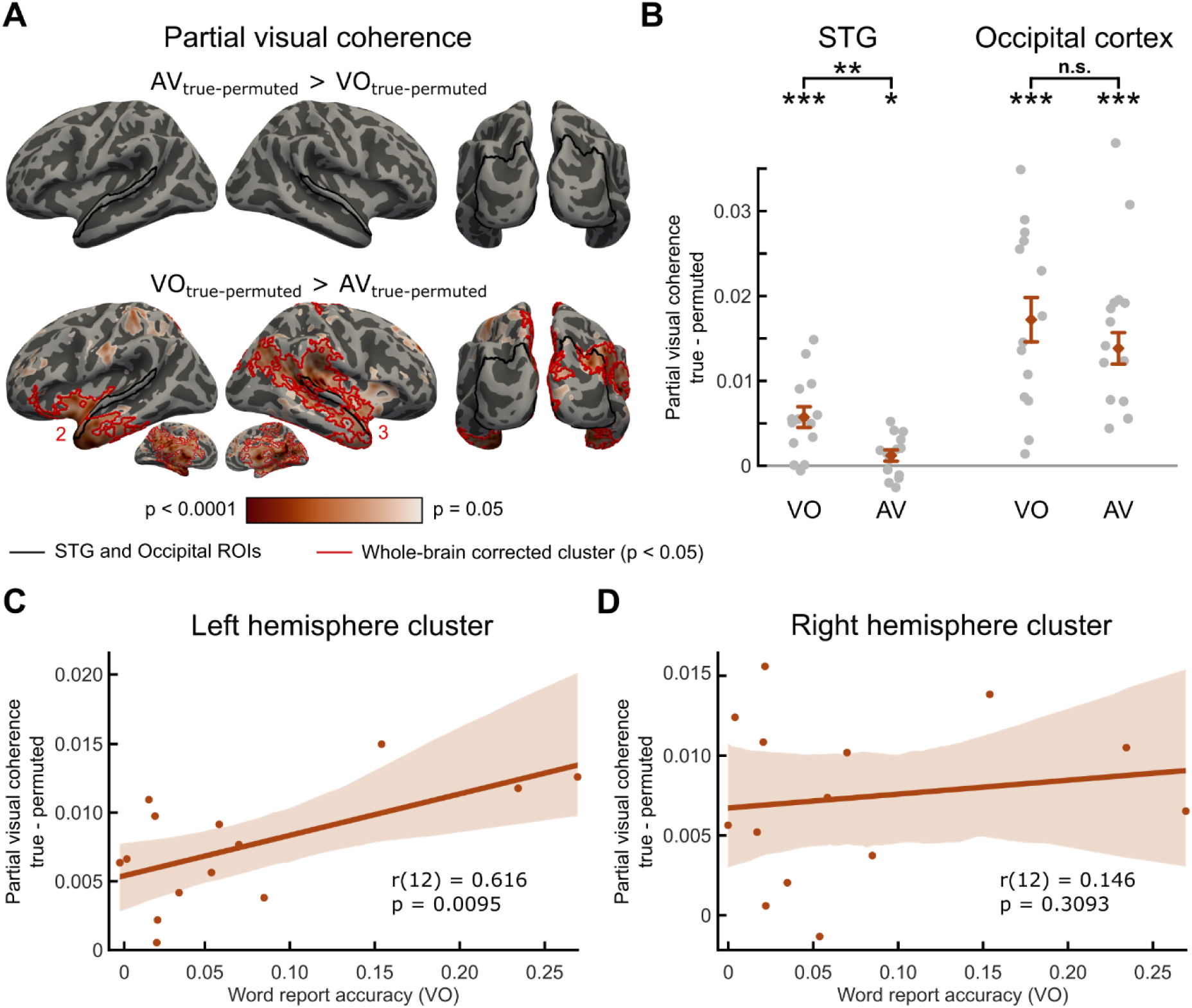
Cross modal influences on visual phase locking in auditory and visual areas. Effect of auditory speech availability on visual phase coherence, whole-brain (A) and ROI-based analysis (B). **A.** Whole-brain maps show significant effects of auditory speech availability (in both directions) on partial visual coherence with respect to permutation-derived null baseline averaged between 2-6 Hz (uncorrected p-values). Black outlines indicate STG and occipital ROIs, red outlines indicate whole-brain corrected clusters (p < 0.05). **B.** Graphs depict the mean auditory partial coherence with respect to the permutation null baseline averaged between 2-6 Hz in AV and VO conditions across ROIs. Diamonds and error bars represent mean ± SEM, grey dots represent individual data points. The clusters supporting the whole-brain corrected significant results are numbered and further details are presented in Table 1. **C** and **D.** The relationship between the effect of auditory speech on visual partial coherence (i.e., VO_true-perm_ - AV_true-perm_) with word report accuracy in VO across participants in the left (C) and right hemisphere (D) clusters (as shown in panel A). Solid lines depict the fitted linear trend, shaded area depicts the 95% CI based on 1000 bootstrap samples. (ROI: region of interest, STG: superior temporal gyrus, AV: audio-visual, VO: visual-only, n.s.: not significant, * p < 0.05, ** p < 0.01, *** p < 0.001)

In occipital cortex (Fig. 6B), we also observed significant partial visual coherence above the permutation baseline both in AV conditions (mean ± SEM = 0.0138 ± 0.0020, t(13) = 6.954, one-tailed p < 0.0001, confirming a finding shown for the full set of trials in Fig. 4B) and VO conditions (mean ± SEM = 0.0173 ± 0.0028, t(13) = 6.171, one-tailed p < 0.0001). However, in contrast to the STG, in occipital cortex there was no evidence that phase-locking to visual speech (partial coherence) was influenced by the presence of auditory speech since VO and AV conditions did not reliably differ (t(13) = -1.978, two-tailed p = 0.0695). However, this difference between STG and occipital cortex ROIs was not confirmed by a two-way interaction since a repeated measures ANOVA comparing partial visual coherence during bimodal and unimodal listening conditions failed to show a significant ROI (STG vs. Occipital cortex) by listening condition (AV vs. VO) interaction (F(1,13) = 0.242, p = 0.6312).

To make sure we are not overlooking other effects through spatially restrictive ROI analyses, we also report equivalent comparisons of bimodal and unimodal speech conditions using whole-brain cluster correction for multiple comparisons. We contrasted whole-brain maps of partial auditory coherence relative to permutation baseline in AV vs. AO conditions and partial visual coherence relative to permutation baseline in AV vs. VO conditions. We found reliable whole-brain corrected effect of visual speech on auditory partial coherence (i.e., AV_true-perm_ > AO_true-perm_) in a right parietal cluster (one-tailed p = 0.0086, cluster 1 in Fig. 5A and Table 1). However, this difference failed to reach corrected significance when intelligibility matched conditions (AV_low_ and AO_high_) were compared (Fig. 5C) and did not correlate with individual differences in our measure of audio-visual benefit (r(12) = -0.151, two-tailed p = 0.6062). Furthermore, supporting ROI results reported in Fig. 6B, we also found reliable whole-brain corrected increase in visual phase-locking for conditions in which auditory speech was absent versus present (i.e., VO_true-perm_ > AV_true-perm_) supported by two large clusters, one in each hemisphere: a left anterior-temporal cluster extending to the midline (one-tailed p = 0.0008, cluster 2 in Fig. 6A and Table 1) and a right temporal and parietal cluster extending to midline structures (one-tailed p = 0.0002, cluster 3 in Fig. 6A and Table 1). This observation confirms that visual speech signals during silent lip reading lead to increased phase-locking in auditory and other language-related brain areas. Furthermore, this effect positively correlates with individual differences in word report for silent speech (lip reading) in the left hemisphere (r(12) = 0.616, one-tailed p = 0.0095, Fig. 6C), but not in the right hemisphere(r(12) = 0.146, one-tailed p = 0.3093, Fig. 6D).

For completeness, we also examined the effect of acoustic clarity (high vs. low) on phase locking to auditory and visual speech in all conditions containing auditory speech signals (i.e., AO_high_, AO_low_, AV_high_ and AV_low_). Neither whole brain, nor ROI-based analysis in occipital cortex and STG identified reliable effects of acoustic clarity on phase locking.

## Discussion

Viewing the face of a conversation partner greatly improves speech comprehension especially under adverse listening conditions (Sumby and Pollack, 1954; Erber, 1975; Summerfield et al., 1992). We provide MEG evidence that both visual and auditory speech signals are tracked by brain responses during speech comprehension (in line with predictions from Schroeder et al., 2008; Peelle and Sommers, 2015) and link these phase-locked neural responses to perceptual outcomes. Both increased acoustic clarity and the presence of visual speech improved behavioural measures of speech comprehension. We also found individual differences in the extent to which listeners benefitted from visual speech compared to clearer auditory speech. This measure of audio-visual benefit correlated with participants’ lip reading ability (Fig. 2C and D). These findings are in line with other results supporting a link between individual differences in lipreading ability and audio-visual benefit in speech comprehension (MacLeod & Summerfield, 1987). However, comparisons of phase-locked MEG responses during silent lipreading and audio-visual benefit show differences between auditory and visual responses which might imply distinct neural mechanisms supporting these two uses of visual speech.

Analysis of coherence between MEG and speech signals identified bilateral and posterior sensors phase-locked to auditory and visual speech, respectively (Fig. 3A). Whole-brain analysis of source-localized MEG responses revealed that auditory speech entrained bilateral temporal, parietal, and inferior frontal areas, while visual speech entrained bilateral occipital areas (Fig. 3B). For audio-visual speech, these areas were entrained more strongly by their respective speech modality (i.e., temporal and parietal areas by auditory speech; occipital areas by visual speech), even when phase-locking to the other speech signal was partialled out (Fig. 4A). These results are comparable with the basic entrainment effects observed in previous studies (Luo et al., 2010; Park et al., 2016; Micheli et al., 2020). Given that auditory and visual speech signals convey correlated information (see Fig 2A, 2B and Chandrasekaran et al., 2009) we used partial coherence analysis to show cross-modal speech processing in auditory and visual sensory areas. Based on previous neurophysiological and brain imaging studies, the most consistently identified sensory regions are STG for auditory speech and occipital cortex (including lateral extrastriate areas) for visual speech (see Beauchamp, 2016 for a review). In the following paragraphs we focus on entrainment effects observed in these two ROIs and consider the nature of the cross-modal influences shown in auditory and visual regions respectively.

### Entrainment to auditory signals in visual cortex

When responding to audio-visual speech, visual cortex reliably tracked both visual and auditory speech signals (Fig. 4B). Although greater phase-locking is observed for visual than for auditory signals (Fig. 4A, B), we also observed significant phase coherence between visual cortex and auditory speech in audio-visual speech conditions (Fig. 4B) and in the absence of visual speech (Fig. 5B and D). A recent study using electrocorticography similarly demonstrated that medial occipital cortex exhibits reliable auditory envelope tracking in the absence of visual speech (Micheli et al., 2020). Other studies have suggested that visual cortex represents unheard auditory speech during silent lipreading by tracking its amplitude envelope (Hauswald et al., 2018) and higher-level linguistic feature representations (Nidiffer et al., 2021; Suess et al., 2022). Correspondingly, we also found evidence of visual cortex tracking the unheard auditory speech envelope in silent lipreading (Fig. 4C). These findings suggest that visual cortices contribute to processing of auditory speech even in normally-hearing and sighted participants (for relevant evidence from blind individuals see Bedny et al., 2011; Watkins et al., 2013).

Critically, however, we also found that auditory speech tracking in the occipital cortex was significantly stronger in the presence of visual speech (i.e., AV conditions) compared to when only auditory speech was available (i.e., AO conditions, see Fig. 5B). This effect has previously been reported in auditory cortex in cocktail party listening (Zion Golumbic et al., 2013). Here, we demonstrate that similar effects can be observed when only a single speaker is present. Furthermore, we show the same influence of visual speech when audio-visual and audio-only conditions are matched for intelligibility (Fig. 2C, 5D). These findings therefore support the hypothesis that visual speech signals enable better phase tracking of auditory speech (Schroeder et al., 2008; Peelle and Sommers, 2015). However, in our study this effect was primarily observed in occipital areas that are not traditionally assumed to make a key contribution to auditory speech perception.

### Entrainment to visual signals in auditory cortex

We consistently observed phase-locking of STG regions to auditory speech signals when listening to audio-visual and audio-only speech (Fig. 3B, 4A), replicating several previous findings (Luo et al., 2010; Peelle et al., 2013; Zion Golumbic et al., 2013; Micheli et al., 2020). In contrast, however, to the bimodal speech processing we reported for visual cortex, we did not find evidence of above-chance phase-locking to visual speech in STG even if both auditory and visual speech signals were available (i.e., AV conditions, Fig. 4B). This might suggest that previous observations of visual phase-locking in auditory brain regions (e.g. Zion Golumbic et al., 2013) are specific to cocktail party listening in which selecting between competing sound sources is required. Importantly, in our study, speech perception was enhanced by the presence of visual speech even when only a single speaker was present. Hence visual enhancement of auditory entrainment in the STG might not be so consistently associated with audio-visual speech processing.

In contrast, when only visual speech was available (VO condition), auditory regions (STG) reliably tracked the visual speech envelope (Fig. 6A, B) and in left hemisphere regions visual entrainment correlated with word report (Fig. 6C). These findings are despite responses to the auditory envelope being absent in visual only conditions (Fig. 4C) and auditory envelope signals being partialled out in analyses of MEG responses (Fig. 6). These findings thus support an account in which the STG only processes visual speech signals when auditory speech information is absent or unavailable. Other studies have demonstrated a similar “fill-in” mechanism in the form of neural reinstatement of noise-masked speech segments (Leonard et al., 2016; Cervantes Constantino and Simon, 2018). Top-down modulation from dorsal stream areas, including motor-related regions have also been proposed to play a role in this visual to phonological mapping (Park et al., 2016; Hauswald et al., 2018). However, in our work, we did not observe differential entrainment of motor regions during silent lip-reading compared to audio-visual speech perception.

### Cross-modal prediction of audio and visual speech signals

Our results show that despite the parallels between silent lip reading and visual benefit to degraded speech perception, distinct neural effects are observed in these two listening situations. Visual cortex shows reliable phase-locking to auditory speech, and this auditory entrainment is enhanced during visual speech processing for audio-visual benefit, or lip-reading (i.e., with and without auditory input). In auditory cortex, however, visual speech does not produce reliable phase-locking, and the presence of visual speech in AV conditions doesn’t significantly enhance phase-locking to auditory speech signals relative to audio-only conditions. Yet, we observed reliable phase-locking to visual speech signals in auditory cortex during silent lip-reading. Thus, despite the behavioural association between silent lip-reading and audio-visual benefit we see marked differences between the influence of visual speech on phase-locked neural responses in auditory and visual cortices. We here offer some tentative suggestions for the interpretation of these findings in line with predictive accounts of speech processing (Hovsepyan et al., 2020; Sohoglu and Davis, 2020) and that have been applied to audio-visual speech perception (Olasagasti et al., 2015).

One interpretation of the present findings is that neural phase-locking to envelope signals provides a timing template such that visual (lip aperture) and auditory (formant frequencies) sensory signals are combined (Olasagasti et al., 2015). Indeed, the phase of cortical theta oscillations in posterior temporal and occipital cortex have been shown to determine whether auditory or visual speech cues determine perception (Thézé et al., 2020), and pre-stimulus oscillatory phase (plausibly determined from visual speech) contributes to identification of ambiguous speech sounds (ten Oever and Sack, 2015). These cross-modal influences can be explained by proposing that visual speech permits predictions for the spectro-temporal properties (i.e., timing and formant frequency) of upcoming speech sounds (van Wassenhove et al., 2005; Arnal et al., 2011) and vice-versa with auditory speech predicting visual signals (Lee and Noppeney, 2014). Auditory prediction errors (and hence auditory phase-locking) arise in visual cortical areas when degraded speech sounds cannot accurately predict visual speech cues during audio-visual speech perception. The resulting prediction errors signal viseme information (Nidiffer et al., 2021) that can be used to update higher-level interpretations and support optimal speech perception when visual and auditory stimuli must be combined (Olasagasti et al., 2015).

When auditory speech is absent (i.e., during silent lip-reading), we continue to observe auditory entrainment in visual cortex. Furthermore, we observe visual envelope cues producing entrainment of auditory brain regions. We interpret this latter finding as also arising from cross-modal predictive processes; in this case due to visually-derived predictions for expected auditory stimuli that are absent. Visually-driven prediction errors expressed in auditory regions encode the absence of expected speech sounds (Blank and von Kriegstein, 2013) and hence drive phase locking in auditory regions. As in other situations in which speech sounds are missing or masked, these prediction errors can reinstate auditory signals and support speech perception (Leonard et al., 2016; Cervantes Constantino and Simon, 2018). Thus, cross-modal prediction errors plausibly explain the pattern of auditory and visual phase-locking observed in two different situations in which visual speech supports word report.

## Acknowledgements

This work was supported by MRC funding of the Cognition and Brain Sciences Unit, supporting MA, LJM, HB, MHD (SUAG/044 G101400). HSØ was supported by a PhD award from the Cambridge Trusts. We would like to thank Clare Cook and Saskia Helbling for their assistance with data collection, Gemma Crickmore and Simon Strangeways for their assistance in recording and editing the speech stimuli.

## Author Contributions

Conceptualization: HSØ, HB, and MHD; Methodology: HSØ and MHD; Software: MA, LJM and HSØ; Formal Analysis: MA, HSØ and MHD; Investigation: HSØ, MHD; Resources: MHD; Data Curation: MA and HSØ; Writing - Original Draft: MA and MHD; Writing - Review & Editing: MA, HSØ, LJM, HB and MHD; Visualization: MA and MHD; Supervision: MHD; Funding Acquisition: MHD, HSØ.

## References

Arnal LH, Wyart V, Giraud A-L (2011) Transitions in neural oscillations reflect prediction errors generated in audiovisual speech. Nature Neuroscience 14:797–801.

Beauchamp MS (2016) Audiovisual Speech Integration: Neural Substrates and Behavior. In: Neurobiology of Language (Hickok G, Small SL, eds), pp 515–526. San Diego: Academic Press. Available at: https://www.sciencedirect.com/science/article/pii/B9780124077942000420 [Accessed June 25, 2021].

Bedny M, Pascual-Leone A, Dodell-Feder D, Fedorenko E, Saxe R (2011) Language processing in the occipital cortex of congenitally blind adults. PNAS 108:4429–4434.

Blank H, von Kriegstein K (2013) Mechanisms of enhancing visual-speech recognition by prior auditory information. Neuroimage 65:109–118.

Bourguignon M, Baart M, Kapnoula EC, Molinaro N (2020) Lip-Reading Enables the Brain to Synthesize Auditory Features of Unknown Silent Speech. J Neurosci 40:1053–1065.

Brainard DH (1997) The Psychophysics Toolbox. Spatial Vision 10:433–436.

Cervantes Constantino F, Simon JZ (2018) Restoration and Efficiency of the Neural Processing of Continuous Speech Are Promoted by Prior Knowledge. Front Syst Neurosci 12 Available at: https://www.frontiersin.org/articles/10.3389/fnsys.2018.00056/full [Accessed June 30, 2021].

Chandrasekaran C, Trubanova A, Stillittano S, Caplier A, Ghazanfar AA (2009) The Natural Statistics of Audiovisual Speech. PLOS Computational Biology 5:e1000436.

Davis MH, Johnsrude IS, Hervais-Adelman A, Taylor K, McGettigan C (2005) Lexical Information Drives Perceptual Learning of Distorted Speech: Evidence From the Comprehension of Noise-Vocoded Sentences. Journal of Experimental Psychology: General 134:222–241.

Destrieux C, Fischl B, Dale A, Halgren E (2010) Automatic parcellation of human cortical gyri and sulci using standard anatomical nomenclature. Neuroimage 53:1–15.

Ding N, Simon JZ (2014) Cortical entrainment to continuous speech: functional roles and interpretations. Front Hum Neurosci 8 Available at: https://www.frontiersin.org/articles/10.3389/fnhum.2014.00311/full [Accessed October 29, 2020].

Erber NP (1975) Auditory-visual perception of speech. J Speech Hear Disord 40:481–492.

Fischl B (2012) FreeSurfer. NeuroImage 62:774–781.

Giraud A-L, Poeppel D (2012) Cortical oscillations and speech processing: emerging computational principles and operations. Nature Neuroscience 15:511–517.

Gramfort A, Luessi M, Larson E, Engemann DA, Strohmeier D, Brodbeck C, Goj R, Jas M, Brooks T, Parkkonen L, Hämäläinen M (2013) MEG and EEG data analysis with MNE-Python. Front Neurosci 7 Available at: https://www.frontiersin.org/articles/10.3389/fnins.2013.00267/full [Accessed April 12, 2021].

Gross J, Hoogenboom N, Thut G, Schyns P, Panzeri S, Belin P, Garrod S (2013) Speech Rhythms and Multiplexed Oscillatory Sensory Coding in the Human Brain. PLOS Biology 11:e1001752.

Gross J, Kujala J, Hämäläinen M, Timmermann L, Schnitzler A, Salmelin R (2001) Dynamic imaging of coherent sources: Studying neural interactions in the human brain. PNAS 98:694–699.

Hartigan JA, Hartigan PM (1985) The Dip Test of Unimodality. The Annals of Statistics 13:70–84.

Hauswald A, Lithari C, Collignon O, Leonardelli E, Weisz N (2018) A Visual Cortical Network for Deriving Phonological Information from Intelligible Lip Movements. Current Biology 28:1453–1459.e3.

Hovsepyan S, Olasagasti I, Giraud A-L (2020) Combining predictive coding and neural oscillations enables online syllable recognition in natural speech. Nat Commun 11:3117.

Hyvarinen A (1999) Fast and robust fixed-point algorithms for independent component analysis. IEEE Transactions on Neural Networks 10:626–634.

Keitel A, Gross J, Kayser C (2018) Perceptually relevant speech tracking in auditory and motor cortex reflects distinct linguistic features. PLOS Biology 16:e2004473.

Kleiner M, Brainard D, Pelli D (2007) What’s new in Psychtoolbox-3? In: Perception, 36 (EVCP Abstract Supplement). Alezzo.

Lakatos P, Shah AS, Knuth KH, Ulbert I, Karmos G, Schroeder CE (2005) An oscillatory hierarchy controlling neuronal excitability and stimulus processing in the auditory cortex. J Neurophysiol 94:1904–1911.

Lee H, Noppeney U (2014) Temporal prediction errors in visual and auditory cortices. Current Biology 24:R309–R310.

Leonard MK, Baud MO, Sjerps MJ, Chang EF (2016) Perceptual restoration of masked speech in human cortex. Nature Communications 7:13619.

Luo H, Liu Z, Poeppel D (2010) Auditory Cortex Tracks Both Auditory and Visual Stimulus Dynamics Using Low-Frequency Neuronal Phase Modulation. PLOS Biology 8:e1000445.

MacLeod A, Summerfield Q (1987) Quantifying the contribution of vision to speech perception in noise. Br J Audiol 21:131–141.

Maris E, Oostenveld R (2007) Nonparametric statistical testing of EEG- and MEG-data. Journal of Neuroscience Methods 164:177–190.

Mattys SL, Davis MH, Bradlow AR, Scott SK (2012) Speech recognition in adverse conditions: A review. Language and Cognitive Processes 27:953–978.

Mégevand P, Mercier MR, Groppe DM, Golumbic EZ, Mesgarani N, Beauchamp MS, Schroeder CE, Mehta AD (2020) Crossmodal Phase Reset and Evoked Responses Provide Complementary Mechanisms for the Influence of Visual Speech in Auditory Cortex. J Neurosci 40:8530–8542.

Micheli C, Schepers IM, Ozker M, Yoshor D, Beauchamp MS, Rieger JW (2020) Electrocorticography reveals continuous auditory and visual speech tracking in temporal and occipital cortex. Eur J Neurosci 51:1364–1376.

Nidiffer AR, Cao CZ, O’Sullivan A, Lalor EC (2021) A linguistic representation in the visual system underlies successful lipreading. bioRxiv:2021.02.09.430299.

Obleser J, Kayser C (2019) Neural Entrainment and Attentional Selection in the Listening Brain. Trends in Cognitive Sciences Available at: http://www.sciencedirect.com/science/article/pii/S1364661319302050 [Accessed October 21, 2019].

Olasagasti I, Bouton S, Giraud A-L (2015) Prediction across sensory modalities: A neurocomputational model of the McGurk effect. Cortex 68:61–75.

Oostenveld R, Fries P, Maris E, Schoffelen J-M (2011) FieldTrip: Open source software for advanced analysis of MEG, EEG, and invasive electrophysiological data. Computational Intelligence and Neuroscience 2011:156869.

O’Sullivan AE, Crosse MJ, Di Liberto GM, Lalor EC (2017) Visual Cortical Entrainment to Motion and Categorical Speech Features during Silent Lipreading. Front Hum Neurosci 10 Available at: https://www.frontiersin.org/articles/10.3389/fnhum.2016.00679/full [Accessed June 23, 2021].

Park H, Kayser C, Thut G, Gross J (2016) Lip movements entrain the observers’ low-frequency brain oscillations to facilitate speech intelligibility King AJ, ed. eLife 5:e14521.

Peelle JE, Davis MH (2012) Neural Oscillations Carry Speech Rhythm through to Comprehension. Front Psychol 3 Available at: https://www.frontiersin.org/articles/10.3389/fpsyg.2012.00320/full [Accessed January 30, 2020].

Peelle JE, Gross J, Davis MH (2013) Phase-locked responses to speech in human auditory cortex are enhanced during comprehension. Cereb Cortex 23:1378–1387.

Peelle JE, Sommers MS (2015) Prediction and constraint in audiovisual speech perception. Cortex 68:169–181.

Rosenberg JR, Halliday DM, Breeze P, Conway BA (1998) Identification of patterns of neuronal connectivity—partial spectra, partial coherence, and neuronal interactions. Journal of Neuroscience Methods 83:57–72.

Schroeder CE, Lakatos P, Kajikawa Y, Partan S, Puce A (2008) Neuronal oscillations and visual amplification of speech. Trends in Cognitive Sciences 12:106–113.

Shannon RV, Zeng F-G, Kamath V, Wygonski J, Ekelid M (1995) Speech Recognition with Primarily Temporal Cues. Science 270:303–304.

Sohoglu E, Davis MH (2020) Rapid computations of spectrotemporal prediction error support perception of degraded speech King AJ, Kok P, Kok P, Press C, Lalor EC, eds. eLife 9:e58077.

Sohoglu E, Peelle JE, Carlyon RP, Davis MH (2014) Top-down influences of written text on perceived clarity of degraded speech. Journal of Experimental Psychology: Human Perception and Performance 40:186–199.

Suess N, Hauswald A, Reisinger P, Rösch S, Keitel A, Weisz N (2022) Cortical Tracking of Formant Modulations Derived from Silently Presented Lip Movements and Its Decline with Age. Cerebral Cortex:bhab518.

Sumby WH, Pollack I (1954) Visual Contribution to Speech Intelligibility in Noise. The Journal of the Acoustical Society of America 26:212–215.

Summerfield Q, Bruce V, Cowey A, Ellis AW, Perrett DI (1992) Lipreading and audio-visual speech perception. Philosophical Transactions of the Royal Society of London Series B: Biological Sciences 335:71–78.

Taulu S, Simola J (2006) Spatiotemporal signal space separation method for rejecting nearby interference in MEG measurements. Phys Med Biol 51:1759–1768.

ten Oever S, Sack AT (2015) Oscillatory phase shapes syllable perception. PNAS 112:15833–15837.

Thézé R, Giraud A-L, Mégevand P (2020) The phase of cortical oscillations determines the perceptual fate of visual cues in naturalistic audiovisual speech. Science Advances 6:eabc6348.

Thomas SM, Jordan TR (2004) Contributions of Oral and Extraoral Facial Movement to Visual and Audiovisual Speech Perception. Journal of Experimental Psychology: Human Perception and Performance 30:873–888.

van Wassenhove V, Grant KW, Poeppel D (2005) Visual speech speeds up the neural processing of auditory speech. PNAS 102:1181–1186.

Voss RF, Clarke J (1975) ‘1/ f noise’ in music and speech. Nature 258:317–318.

Watkins KE, Shakespeare TJ, O’Donoghue MC, Alexander I, Ragge N, Cowey A, Bridge H (2013) Early Auditory Processing in Area V5/MT+ of the Congenitally Blind Brain. J Neurosci 33:18242– 18246.

Yi A, Wong W, Eizenman M (2013) Gaze patterns and audiovisual speech enhancement. J Speech Lang Hear Res 56:471–480.

Zion Golumbic E, Cogan GB, Schroeder CE, Poeppel D (2013) Visual Input Enhances Selective Speech Envelope Tracking in Auditory Cortex at a “Cocktail Party.” J Neurosci 33:1417–1426.

Zoefel B, Allard I, Anil M, Davis MH (2020) Perception of Rhythmic Speech Is Modulated by Focal Bilateral Transcranial Alternating Current Stimulation. Journal of Cognitive Neuroscience 32:226–240.

Zoefel B, Archer-Boyd A, Davis MH (2018) Phase Entrainment of Brain Oscillations Causally Modulates Neural Responses to Intelligible Speech. Current Biology 28:401–408.e5.

